# Dopaminergic drug treatment remediates exaggerated cingulate prediction error responses in obsessive-compulsive disorder

**DOI:** 10.1101/225938

**Authors:** Graham K Murray, Franziska Knolle, Karen D Ersche, Kevin J Craig, Sanja Abbott, Shaila S Shabbir, Naomi A. Fineberg, John Suckling, Barbara J Sahakian, Edward T Bullmore, Trevor W Robbins

**Affiliations:** Department of Psychiatry, University of Cambridge; Behavioural and Clinical Neuroscience Institute, University of Cambridge; Cambridgeshire and Peterborough NHS Foundation Trust; Department of Psychology, University of Cambridge; GlaxoSmithKline, Immuno-Inflammation Therapeutic Area Unit, Stevenage UK; European Bioinformatics Institute; Department of Psychiatry, Queen Elizabeth II Hospital, Welwyn Garden City, United Kingdom

**Keywords:** Obsessive-compulsive disorder, Amisulpride, Pramipexole, reward learning, prediction error, anterior cingulate, nucleus accumbens, computational model

## Abstract

**Rationale:** Patients with obsessive-compulsive disorder (OCD) have been found to show exaggerated error responses and prediction error learning signals in a variety of EEG and fMRI tasks, with data converging on the anterior cingulate cortex as a key locus of dysfunction. Considerable evidence has linked prediction error processing to dopaminergic function.

**Objective:** In this study we investigate potential dopaminergic dysfunction during reward processing in the context of OCD.

**Methods:** We studied OCD patients (n=18) and controls (n=18) whilst they learned probabilistic associations between abstract stimuli and monetary rewards in the fMRI scanner involving administration (on separate visits) of: a dopamine receptor agonist, pramipexole 0.5mg; a dopamine receptor antagonist, amisulpride 400mg, and placebo. We fitted a Q-learning computational model to fMRI prediction error responses; group differences were examined in anterior cingulate and nucleus accumbens regions of interest.

**Results:** There were no significant group, drug or interaction effects in number of correct choices; computational modeling suggested a marginally significant difference in learning rates between groups (p=0.089, partial ⍰^2^=0.1). In the imaging results, there was a significant interaction of group by drug (p=0.013, partial ⍰^2^=0.13). OCD patients showed abnormally strong cingulate signaling of prediction errors during omission of an expected reward, with unexpected reduction by both pramipexole and amisulpride (p=0.014, partial ⍰^2^=0.26, 1-β error probability=0.94). Exaggerated cingulate prediction error signaling to omitted reward in placebo was related to trait subjective difficulty in self-regulating behavior in OCD.

**Conclusions:** Our data support cingulate dysfunction during reward processing in OCD, and bidirectional remediation by dopaminergic modulation, suggesting that exaggerated cingulate error signals in OCD may be of dopaminergic origin. The results help to illuminate the mechanisms through which dopamine receptor antagonists achieve therapeutic benefit in OCD. Further research is needed to disentangle the different functions of dopamine receptor agonists and antagonists during bidirectional modulation of cingulate activation.

## Introduction

Obsessive-compulsive disorder (OCD) has been associated with deficits in learning and decision making (Gillan and Robbins, 2014; Nielen *et al*, 2009). OCD patients were found to show reduced response control in punishment trials, with impulsivity being correlated with severity of symptoms (Morein-Zamir *et al*, 2013), and abnormal error responses in a variety of response conflict and error monitoring tasks using EEG (Gillan *et al*, 2017; Mathews *et al*, 2012), suggesting overactivity of the performance monitoring system involving the anterior cingulate cortex (Hammer *et al*, 2009).

Neural mechanisms underlying decision making and learning involve fronto-striatal networks generating reward prediction errors which are highly sensitive to dopaminergic modulation (Frank *et al*, 2004; Kehagia *et al*, 2010; Schultz and Dickinson, 2000). Extensive evidence from studies in experimental animals has demonstrated that dopamine neurons show increased rates of firing to unexpected or better than expected rewards (positive prediction error), and a reduction in firing rate during omission of expected rewards (negative prediction error) (Schultz *et al*, 1997). Studies in healthy human volunteers have shown dopaminergic modulation of the brain response to a reward prediction error response using fMRI (Bernacer *et al*, 2013; Eisenegger *et al*, 2014; Pessiglione *et al*, 2006). Evidence from human molecular imaging studies of patients with OCD suggests a possible role for dopaminergic pathology in OCD (Denys *et al*, 2004; Hesse *et al*, 2005; Moresco *et al*, 2007; Olver *et al*, 2009, 2010; Perani *et al*, 2008; Sesia *et al*, 2013; Van Der Wee *et al*, 2004). Whilst dopaminergic medications are not first-line treatments for OCD, dopamine receptor antagonists are often used to augment first-line treatment in refractory cases (Hirschtritt *et al*, 2017), and the efficacy of this strategy is confirmed by meta-analyses (Dold *et al*, 2015; Veale *et al*, 2014).

Notably, the nucleus accumbens (part of the ventral striatum) receives dense dopaminergic projections and is a target for deep brain stimulation in OCD (Denys *et al*, 2010). A prior study demonstrated abnormal reward processing in the nucleus accumbens in OCD (Figee *et al*, 2011): it reported attenuated activations for reward anticipation, but did not examine learning or prediction error signaling. The nucleus accumbens has notably been associated with coding prediction error in healthy controls (Abler *et al*, 2006; O’Doherty *et al*, 2004; Pagnoni *et al*, 2002; Rodriguez *et al*, 2006). The concept of negative prediction error is closely related to that of error monitoring, which is reliably abnormal in OCD, although the two processes are not identical. Both processes involve evaluating worse than expected outcome, motivating the study of prediction error signaling in OCD. Recently, Hauser *et al*. (2017) reported increased brain signals in response to reward prediction errors in the anterior cingulate cortex and the putamen, indicating an overactivation of the monitoring system. Furthermore, dopamine dependent activation in the anterior cingulate has also been associated with increased error awareness and performance monitoring (Hester *et al*, 2012; Pauls *et al*, 2012). Reward prediction error processing is closely linked to the dopaminergic system (Schultz *et al*, 1997). A plausible hypothesis unifying (I) the known role of dopamine in signaling reward prediction errors, (II) the existing evidence of dopaminergic pathology in OCD, and (III) evidence of abnormal error processing in OCD, is that OCD patients may exhibit abnormal processing of reward prediction errors, especially during omission of expected reward linked to abnormal dopaminergic function.

To our knowledge, no study has yet examined a direct modulation of the dopaminergic system on a reward prediction error task in OCD. In the current experiment, we therefore compared the effect of dopaminergic drug treatments on prediction error signaling in OCD. We recruited a group of patients with OCD and a group of healthy volunteers as controls, and examined brain responses during studies of reinforcement learning in an fMRI scanner after administration of a dopamine D_2/3_ receptor agonist, pramipexole, or a dopamine D_2/3_ receptor antagonist, amisulpride, or a placebo. We hypothesized that fronto-striatal reward prediction error signaling in OCD would be exaggerated under placebo; given the evidence for dopamine receptor antagonism add-on treatment for OCD, we hypothesized that this abnormality would be normalized by amisulpride but not by pramipexole, which has been shown in some cases to worsen compulsivity (Kolla *et al*, 2010). Our hypotheses were principally focused on two brain regions of particular interest: we predicted that dopaminergic abnormalities in anterior cingulate reward prediction errors would be prominent in OCD during processing of omitted rewards, as this is analogous to the enhanced cingulate error signaling suggested by Error Related Negativity studies in OCD (Gillan *et al*, 2017). A secondary hypothesis was that in the nucleus accumbens (Figee *et al*, 2011) there would be dopaminergic reward prediction error abnormalities in OCD (during reward receipt and/or omission).

## Methods

### Participants

Thirty-six right-handed volunteers participated in this study (Table 1): 18 individuals satisfying DSM-IV-TR criteria for OCD, and 18 healthy controls. The groups were matched for age, gender, IQ and years of education. Prior to study enrolment, all participants had a satisfactory medical review and were screened for any other current Axis I psychiatric disorder using the Structured Clinical Interview for the *DSM-IV-TR* Axis I psychiatric disorders. OCD patients were recruited from a specialist OCD clinic and through independent charities and most were taking serotonin selective reuptake inhibitor (SSRI) medication (see Supplementary Material for details). Participants were excluded based on any other Axis I psychiatric disorders. All volunteers provided written informed consent. The study was approved by the Cambridge Research Ethics Committee.

**Table 1.**
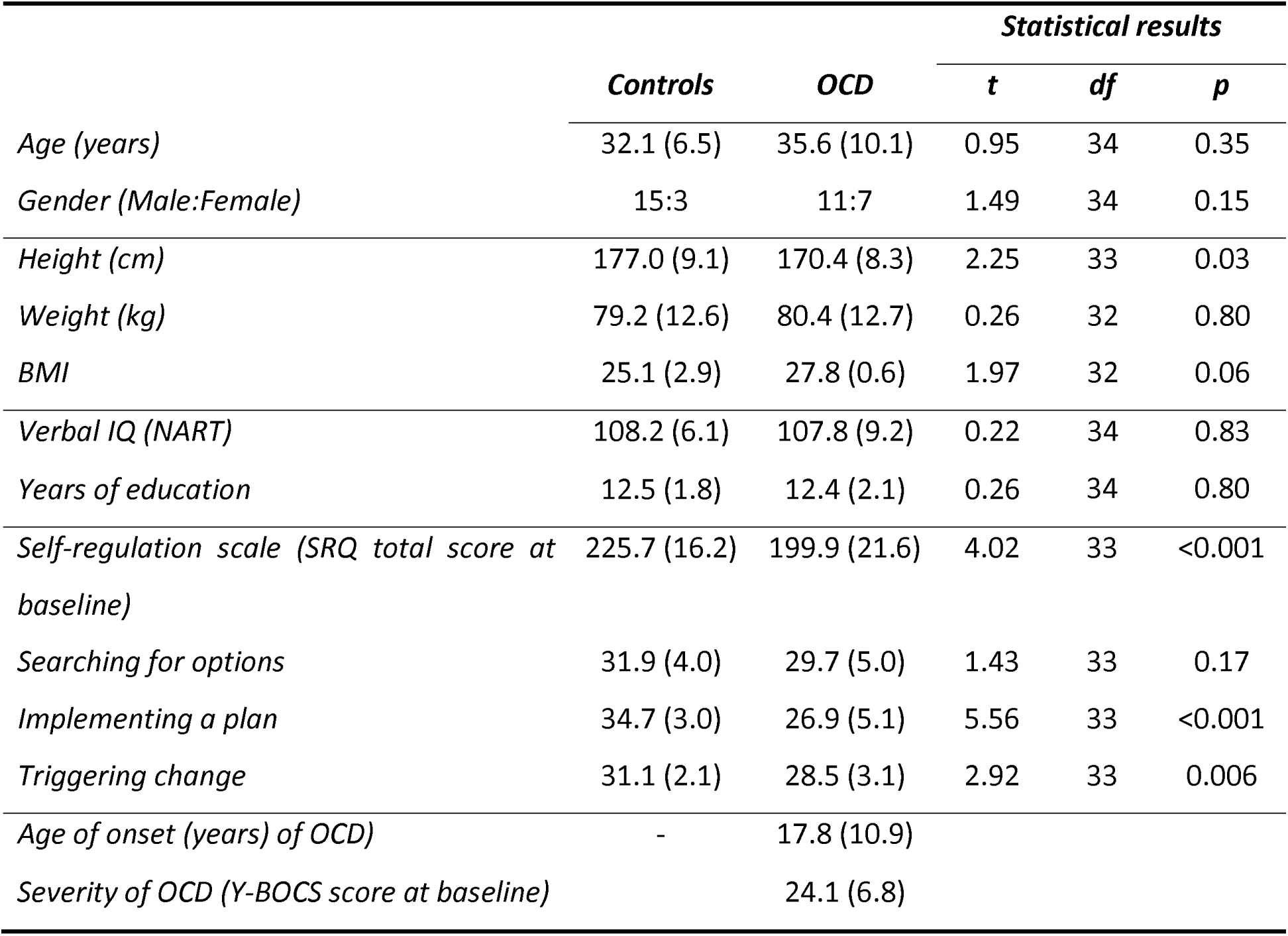
Subject characteristics. Mean scores (SD) are shown for continuously distributed variables according to diagnostic group.

### Pharmacological interventions

Participants attended for three in-unit assessments in which treatments were administered by mouth. On one visit, they received 0.5mg pramipexole, a selective agonist at the dopamine D_2/3_ receptors; on another visit 400mg amisulpride, a selective antagonist at the dopamine D_2/3_ receptors, and on another visit a placebo. Drug administration was conducted in a double-blind, placebo controlled fashion counterbalanced for drug/visit order. Each dosing of drug/placebo was administrated 60min prior to scanning to assure peak plasma levels for both drugs during scanning. The time-point of drug administration was based on pharmacokinetic data for both drugs (Coukell *et al*, 1996; Rosenzweig *et al*, 2002; Wright *et al*, 1997).

Patients’ eligibility to proceed with the scanning was assured by ECG-monitoring. We initially administered a single oral dose of 1.5mg of pramipexole to the first three healthy volunteers. However, this dose of pramipexole was poorly tolerated, the three healthy volunteers were unable to perform the tasks at this treatment session because of nausea, vomiting, sweating, and tiredness. Subsequently, the dose of pramipexole was reduced to 0.5mg orally for all participants (i.e., for 18 controls, including the three volunteers who did not tolerate the higher dose). All participants were also administered a total of 30mg of domperidone orally for each treatment session to prevent emetic effects of dopamine receptor agonism. The administration was split up over three time-points. 10mg of domperidone was to be taken 12 hours and 2 hours before arrival, and further 10 mg of domperidone were administered together with the study medication.

Subjective drug effects were assessed at two time-points, 1 and 1.5 hours after drug administration using the Bond-Lader Visual Analogue Scale (Bond and Lader, 1974). The time-points referred to assessment immediately before and after fMRI scanning. At this time-points blood-samples were taken from the participants, to assess plasma-levels and prolactin concentration.

### fMRI task

During the fMRI scan, subjects carried out a probabilistic learning task that required making choices to maximize wins and minimize losses (Figure 1), adapted from previous similar tasks (Bernacer *et al*, 2013; Ermakova *et al*, 2018; O’doherty *et al*, 2004; Pessiglione *et al*, 2006). In each trial, one of three possible pairs of abstract pictures was randomly presented: rewarding, punishing, or neutral (40 trials of each valence). For each trial, the subject used a button push to indicate a choice of picture. When viewing the potentially rewarding pair, selection of one of the pictures led to a financial win with a 70% probability and of a no-change outcome with 30% probability, whereas the selection of the other picture led to gain with only 30% probability. The potentially punishing pair led to a financial loss on 70% and 30% trials depending on stimulus choice, and the neutral pair led to no change. We focus in this report on how participants learned about rewards; hence we report fMRI results from the reward trials only. Details of behavior on other trial types are reported in the Supplementary Material.

**Figure 1.**
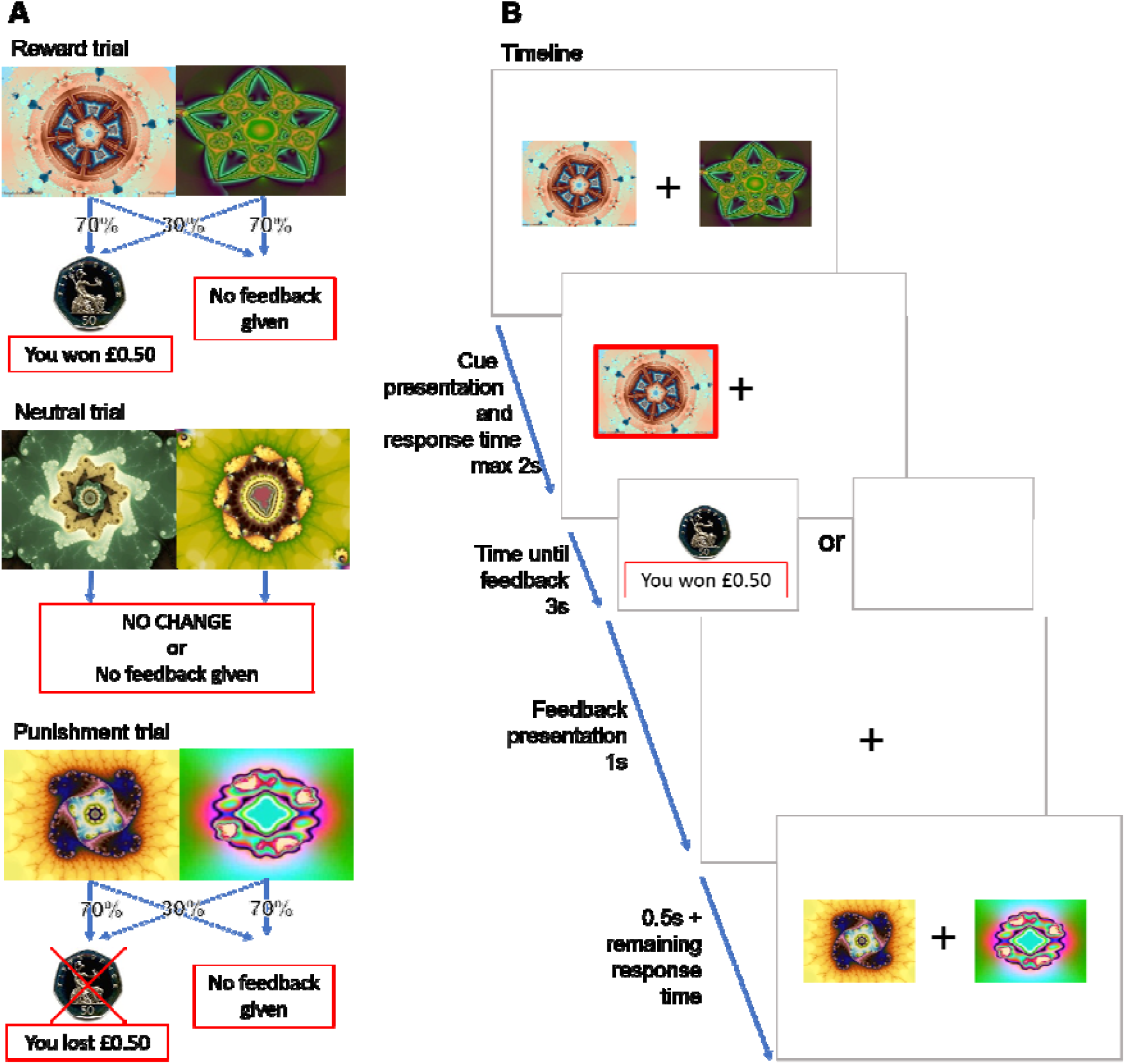
Panel A presents the three different trial types and feedback probabilities. Panel B presents the experimental task, including trial timing. With cue presentation participants have a maximum of 2s to make a decision. The chosen stimulus is circled in red and presented for 3s. Feedback or no feedback is presented for 1s, which is followed by a fixation cross for a minimum of 0.5s (0.5s + (2s – reaction time)).

### Behavioral analysis

We used a repeated measure analysis of variance (ANOVA) to investigate the group differences (controls, OCD; between subjects’ variable) and effect of drug (placebo, pramipexole, amisulpride; within subjects’ variable) in stimuli choices and reaction times. We were interested in the proportion with which the subjects gave a ‘correct’ response and whether this performance was dependent on trial types and drugs. Stimulus selection was considered correct when choosing the stimulus with the high-probability of leading to winning of 50 pence in the reward trial (regardless of whether or not the participant was rewarded on that particular trial), the picture with the high probability of receiving a neutral feedback picture in neutral trials, and stimulus with the high-probability of leading to no punishment. In the neutral trials, the assignment of ‘correctness’ is arbitrary, but assigning one stimulus in each category to be the correct stimulus allows examination of to what extent response patterns differed across trial types.

Computational learning model parameters (see below) were analyzed in a repeated measures ANOVA with group as a between subjects’ variable and drug as a within-subjects’ variable.

### Computational modeling of learning

We estimated the positive and negative reward prediction error value for each trial, based on a basic Q learning algorithm (Murray *et al*, 2008; Pessiglione *et al*, 2006), that involves modelling learning rates (alpha) and “exploration-exploitation” parameters (betas). Beta, also understood as the inverse temperature, balances between random sampling or exploration of actions, and the exploitation of current knowledge. See the Supplementary Material for detailed description of the computational modelling methods.

### MRI analysis

MRI acquisition is described in the Supplementary Material. Data were initially analyzed a voxelwise fashion using FSL software (Jenkinson *et al*, 2012). Data were analyzed using the FSL tool, FEAT (FMRI Expert Analysis Tool) using the following steps. EPI images were realigned, motion-corrected, and spatially smoothed with a Gaussian kernel of 6mm (full-width half-maximum). The time series in each session was high-pass filtered (60s cut off), and the images were registered to the structural MPRAGE scan obtained from the corresponding subject (having first removed the skull using the brain extraction tool BET), and finally transformed into standard space using the MNI template.

In the analysis of fMRI data, eight explanatory variables (EVs), along with their temporal derivatives (used instead of slice timing correction), were defined in our model as follows: 1) Reward expected value; 2) Positive reward prediction error; 3) Negative reward prediction error; 4) Punishment expected value 5) Positive Punishment PE; 6) Negative Punishment PE; 7) Feedback presentation; 8) Cue presentation. FSL requires all events to be set with non-zero duration: we set all to be 2sec. EVs1-6 were based on the trial by trial computational model estimates of reward cue values and prediction errors. The timings of EVs1 and 4 correspond to cue presentation. EVs2,3,5 and 6 correspond to the times of delivery of reward or omission of expected reward. EV2 represents reward prediction error when a reward is received (i.e. positive prediction error), and EV3 represents the prediction error when an expected reward is omitted (negative prediction error). A positive EV3 parameter estimate, therefore, reflects lower BOLD values during omission of predicted rewards. Positive and negative reward prediction error activity were considered separately, as prior research in OCD has focused on brain correlates of errors using tasks where an error is usually accompanied by negative feedback or represents some kind of mistake, which is more analogous to negative prediction error than positive prediction error. In the present analysis, we focused on EV2 and EV3, the positive and negative reward prediction errors. We computed separate +1 contrasts on EV2, and on EV3 at the single subject level (contrast of parameter estimates, or COPEs). We defined two regions of interest (ROIs: see below and Supplementary Material) and extracted average COPE values from each participant in these ROIs, in order to subject these to ANOVA in the statistics package SPSS. Before ANOVA was performed, we examined the distributions of the COPE values and excluded outliers more than three standard deviations away from the mean of all the samples pooled (Howell, 2009). Although psychologically, punishment learning is of considerable interest in OCD, the neurochemical basis of punishment prediction error signals is much less clear than those of reward prediction errors, with prior studies (Pessiglione *et al*, 2006) failing to find evidence of dopaminergic modulation of punishment prediction error learning signals. For this reason, although our paradigm included punishment learning, we did not proceed to formulate or test hypotheses regarding punishment brain signals.

In this study, we were interested in the anterior cingulate cortex, using a sphere, 14mm, centered at x=0, y=42, z=18, based on (Hauser *et al*, 2017), and the nucleus accumbens, as defined as in the Harvard-Oxford-subcortical atlas supplied with FSL, during prediction error processing.

### Relation to behavioral scales

We determined relationships between anterior cingulate and accumbens prediction error COPE values in placebo and behavioral scales using nonparametric correlations (Spearman’s rho). We examined total OCD symptoms (in patients only) as measured by the Y-BOCS scale (Goodman *et al*, 1989) and a subjective assessment of goal directed behavior/cognitive control (in both groups) using the self-regulation questionnaire (SRQ), which assesses the trait ability to develop, implement and flexibly maintain planned behavior in order to achieve one’s goals (Brown *et al*, 1999). It contains items for receiving relevant information, evaluation, triggering change, searching for option, formulating a plan, implementing a plan and assessing effectiveness.

## Results

### Discrimination learning

Independent of drug, participants learned to choose the picture that most often led to winning 50p (high-probability stimulus: correct response) in the majority of reward trials (mean percentage of correct choices for controls: 83.58% (±26.3) and patients: 72.45% (±34.9)); there was a tendency for OCD patients to make more errors across all treatment conditions (Figure 2) (marginal effect: F=3.5, dfd=1, dfe= 34, p=0.070, partial ⍰^2^=0.09), but no effect of drug (p=0.39) or group X drug interaction (p=0.86). (See Supplementary Material for learning curves for reward trials, and description of other trial types.)

**Figure 2.**
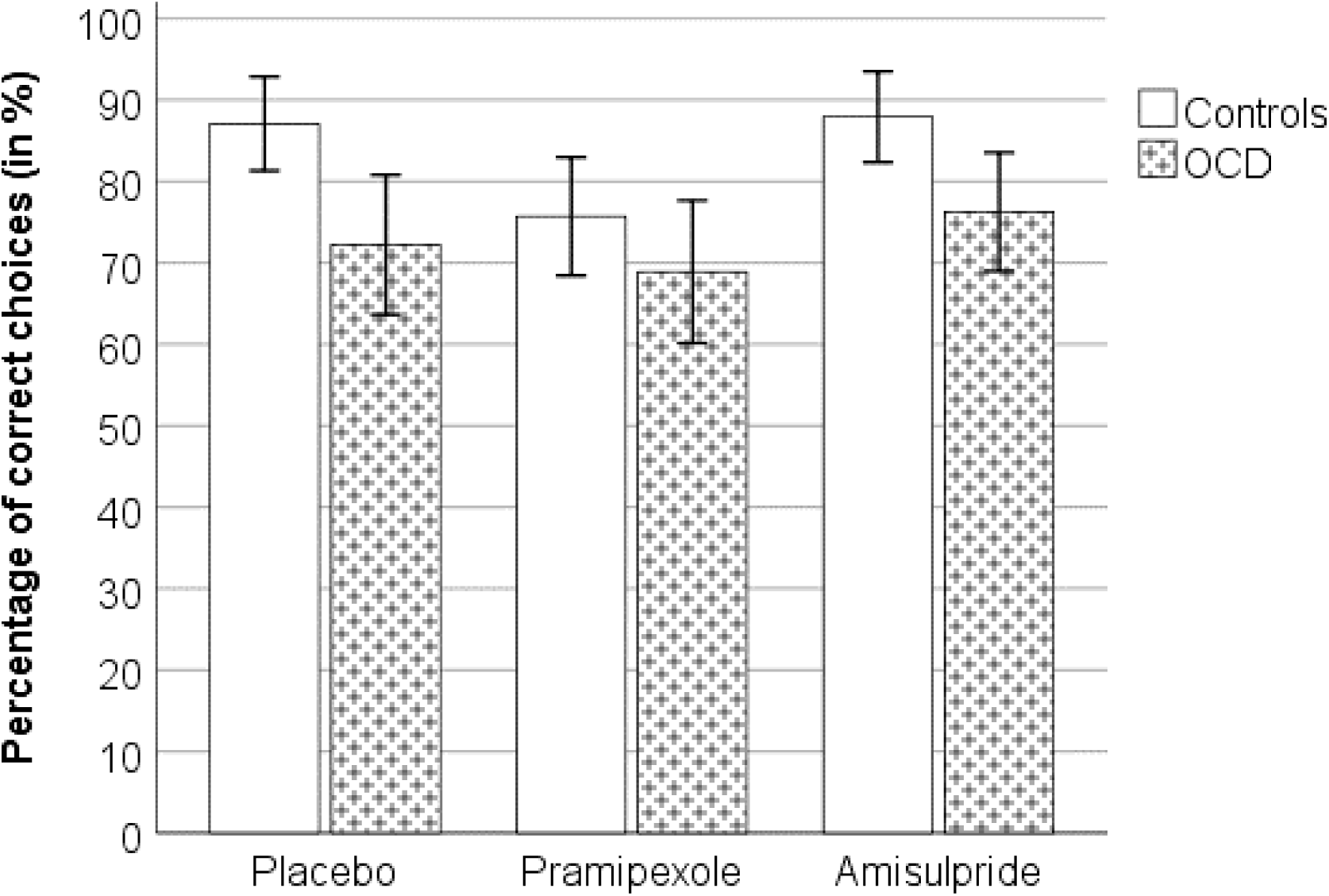
Choice performance (mean ±1 SEM) during reward trials under different drugs, separated by group.

### Reaction time results

There was a significant main effect of drug (F=5.63, dfd=2, dfe=68, p=0.005, partial ⍰^2^=0.14), but no effect of group (p=0.62) or an interaction (p=0.70). Bonferroni corrected post-hoc tests revealed that participants independently of group reacted to the reward trials significantly slower under pramipexole (845.29ms ±29.22) compared to placebo (770.93ms ±23.66): p=0.016, or amisulpride (757.86ms ±26.41): p=0.002 (see Supplementary Material Results and Table 1 for more details).

### Computational modelling parameters

For the learning rate, *α*, we found a marginal effect of group (F=3.07, dfd=1, dfe=34, p=0.089, partial ⍰^2^=0.1), but no effect of drug (p=0.94) or an interaction (p=0.31). The marginally significant group effect suggests that controls, independent of drug, have a higher learning rate (mean α=0.27, Standard Error 0.02) than OCD patients (mean *α*=0.22, standard error 0.02). For β, there were no differences across groups (p=0.82), drug (p=0.1) or an interaction (p=0.78). There was no effect of drug (p=0.38), group (p=0.22) or interaction (p=0.54) on model fit.

### Imaging results, pooled data (“main effect of task”)

Results from pooled data, collapsing over group and drug conditions, are presented in Supplementary Material.

### Imaging results: Anterior cingulate cortex, negative prediction error

The mean COPE values for negative prediction error in the anterior cingulate cortex were extracted to enable a region of interest analysis in accordance with our hypothesis. Data from three OCD patients were excluded because, for at least one of the sessions, the mean COPE value from the ROI was over three standard deviations from the mean of the entire sample. Following repeated measure ANOVA, there were no main effects of group (p=0.88) or drug (p=0.44). However, there was a significant interaction of group by drug (F=4.65, dfd=1,2, dfe=30,62, p=0.013, partial ⍰^2^=0.13, 1-β error probability=0.7). Within each group, we then conducted a Bonferroni corrected post hoc test analyzing the effect of the drug with a repeated measure ANOVA. In controls, the drug intervention did not influence negative reward prediction error processing, as shown by a non-significant drug effect (p=0.16). In the OCD patients, however, there was a significant drug effect (F=5.01, dfd=2, dfe=28, p=0.014, partial ⍰^2^=0.26, 1-β error probability=0.94). On further analysis within the patient group, amisulpride (mean=−40.03, SD=157.35) as well as pramipexole (mean=−20.83, SD=133.09) led to significant differences compared with placebo but not between themselves (mean=128.56, SD=189.54): placebo > amisulpride, p=0.030; placebo > pramipexole, p=0.017; amisulpride > pramipexole, p=0.70.

Furthermore, we applied a univariate ANOVA to each drug intervention separately (Figure 3). For placebo, there was a significant effect for group (F=8.84, dfd=1, dfe=31, p=0.006, partial ⍰^2^=0.22, 1-β error probability=0.91), showing that the COPE values in the controls (mean=−42.21, SD=140.24) were lower than in OCD (mean=128.57, SD=189.54). For the other two interventions, no significant differences were found between groups (amisulpride: controls (mean=15.12, SD=243.59) > OCD (mean=−40.03, SD=157.35): p=0.46; pramipexole: controls (mean=117.4, SD=362.97) > OCD (mean=−20.83, SD=133.09): p=0.17).

**Figure 3.**
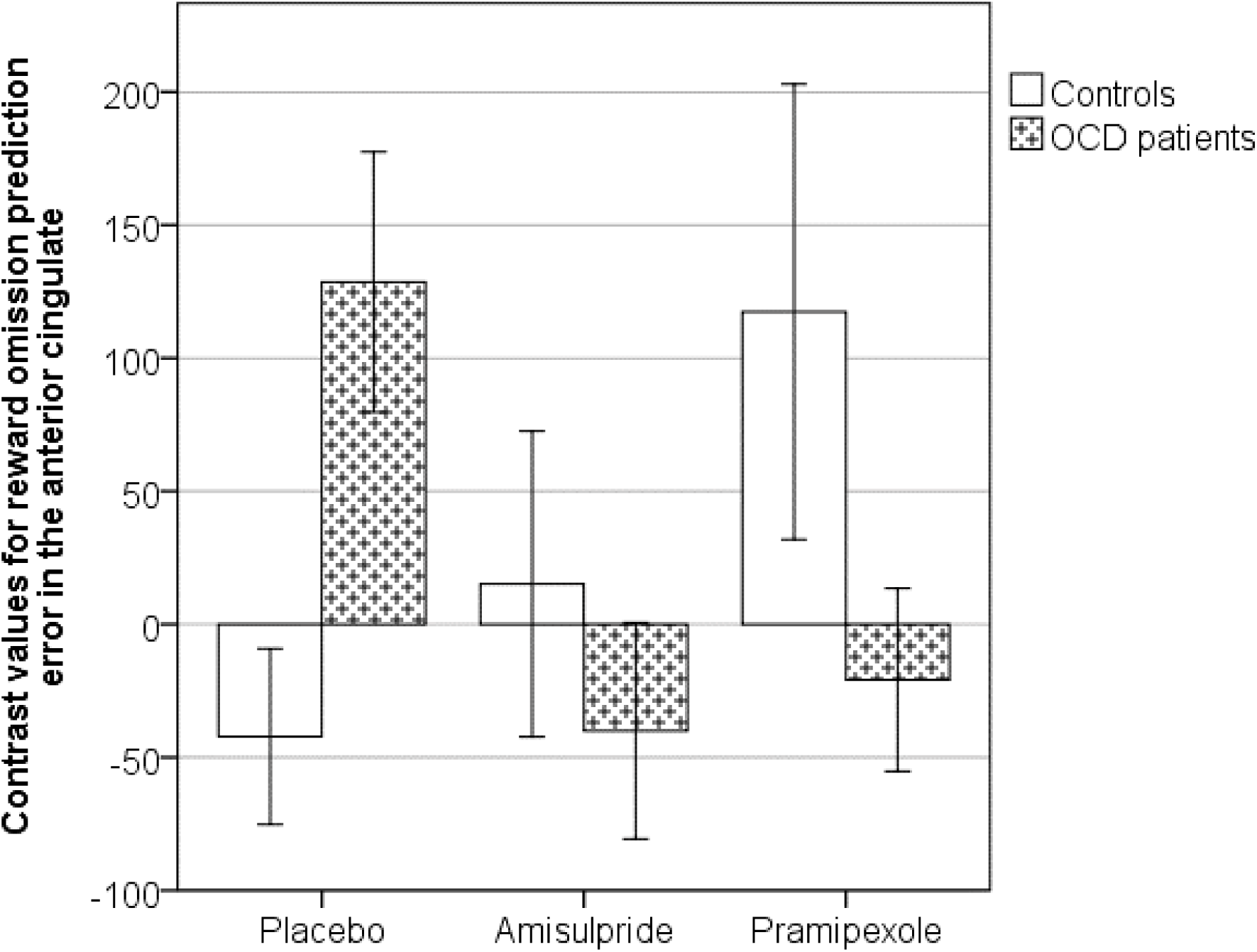
Bar chart showing mean negative reward prediction error contrast value (±1 SEM) extracted from the anterior cingulate cortex at the time of reward omission. A higher contrast value indicates that the BOLD signal was positively correlated with negative prediction error. OCD patients were significantly different from controls following placebo, but both groups behave similarly following amisulpride or pramipexole.

### Imaging results: Anterior cingulate cortex, positive prediction error

There were no significant group, drug or interaction effects for positive prediction error in the anterior cingulate cortex in a repeated measure ANOVA.

### Imaging results: Nucleus accumbens, negative prediction error

There were no significant group or drug or interaction effects on negative prediction error COPE values in the nucleus accumbens.

### Imaging results: Nucleus accumbens, positive prediction error

The mean COPE values for positive prediction error in the nucleus accumbens were extracted to enable a region of interest analysis in accordance with our hypothesis. Data from three controls and one OCD patient were excluded as on at least one of the different treatment sessions the mean COPE values from the ROI were over three standard deviations from the mean of the entire sample. Following repeated measure ANOVA, there was a significant main effect of group (F=11.82, dfd=1, dfe=30, p=0.002, partial ⍰^2^=0.28, 1-β error probability=0.96), but no significant effect for drug (p=0.45) or interaction of group by drug (p=0.54).

We then applied a univariate ANOVA to each drug intervention separately (Figure 4). For placebo, there was a significant effect for group (F=5.11, dfd=1, dfe=30 p=0.031, partial ⍰^2^=0.15, 1-β error probability=0.75), showing that the COPE values in the controls (mean=32.28, SD=120.46) were significantly less than in OCD group (mean=125.74, SD=113.32). For the amisulpride intervention, no significant differences were found between groups (amisulpride: controls (mean=89.15, SD=125.77) > OCD (mean=110.81, SD=122.66): p=0.63). For the pramipexole intervention, we found a marginal effect (F=2.90, dfd=1, dfe=30, p=0.099, partial ⍰^2^=0.09) with controls (mean=23.36, SD=124.58) showing less activity compared to OCD patients (mean=93.21, SD=107.35).

**Figure 4.**
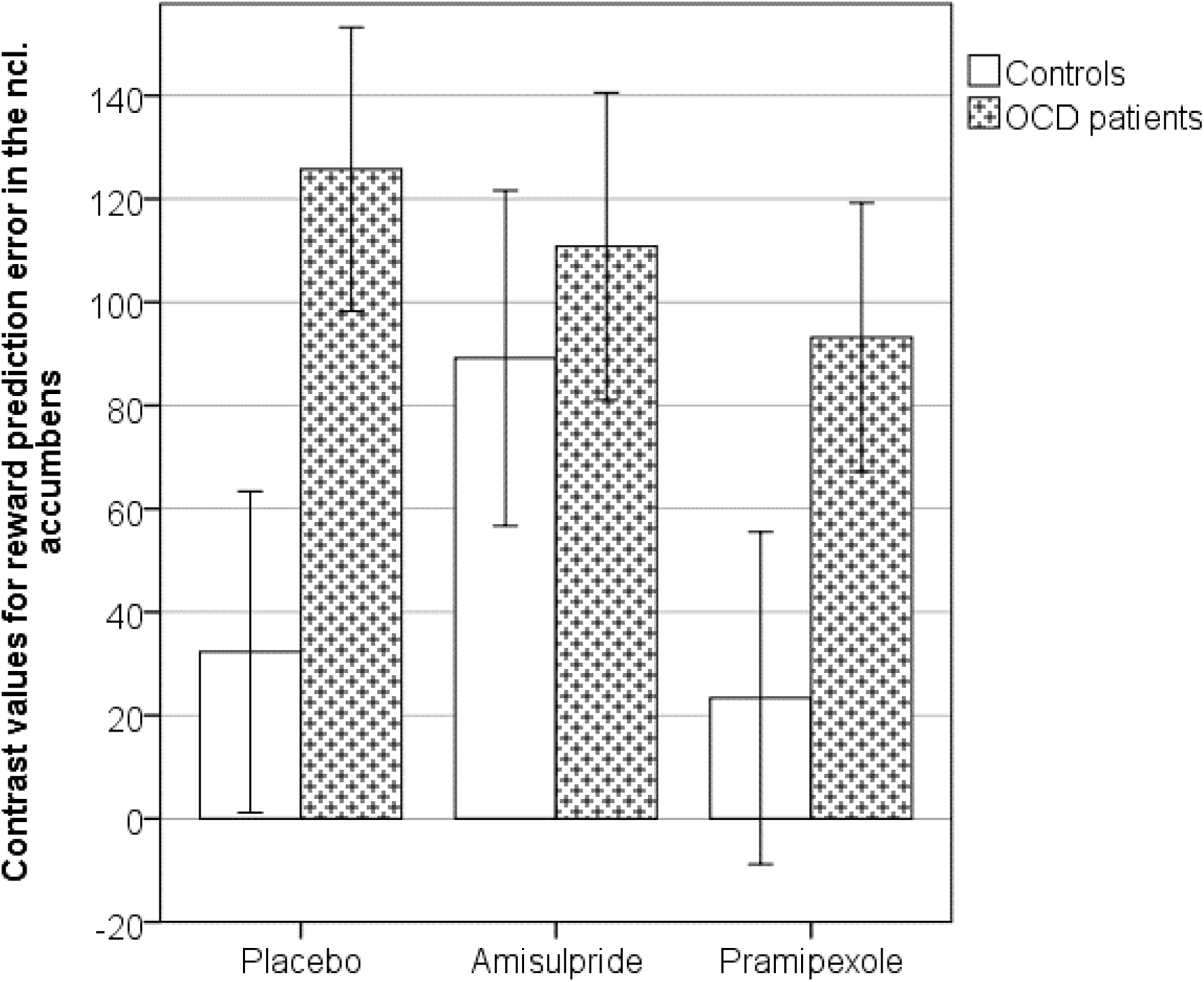
Bar chart showing mean reward prediction error contrast values (±1 SEM) extracted from the nucleus accumbens. OCD patients were significantly different from controls following placebo, but both groups behave similarly following amisulpride or pramipexole.

We also conducted a Bonferroni corrected post hoc test analyzing the effect of the drug within each group using repeated measure ANOVA. Neither in controls, nor in OCD patients, did the drug intervention influence reward positive prediction error processing (controls: p=0.99, OCD patients: p=0.60).

### SSRI Treatment – medication interaction

In order to control for a potential interaction of the patients’ individual routine medication and the study drug treatment on reward prediction error signaling, we created a dichotomized medication variable (high dose or not high dose, the latter including two medication free patients) and used this variable as a between-subject factor in a repeated measure ANOVA on the three treatment types (i.e. placebo, pramipexole, amisulpiride). We found a significant treatment effect on anterior cingulate cortex negative prediction error COPE values (F=5.42, dfd=2, dfe=6, p=0.011), as in our analysis above without the medication factor, but no SSRI medication dose effect (p=0.8) or interaction (p=0.3). In the nucleus accumbens, we did not find any significant effect of high SSRI dose or interaction on positive prediction error COPE values (all p>0.1).

### Correlations with behavioral scales

There was no correlation between placebo cingulate negative prediction error COPE values and OCD symptoms as measured by Y-BOCS score (rho=−0.2, p=0.4). However, there was a significant negative correlation between the cingulate negative prediction error signal and trait self-regulation as measured by the SRQ (rho=−0.7, p=0.002, including outliers; rho=−0.75, p=0.002, excluding outliers), such that patients who reported more difficulty with self-regulation also demonstrated exaggerated negative error signals in the anterior cingulate (Figure 5). On analyses of subscales, the overall association was mainly driven by associations with subscales for “triggering change”, “searching for options” and “implementing a plan” (all p<0.005).

**Figure 5.**
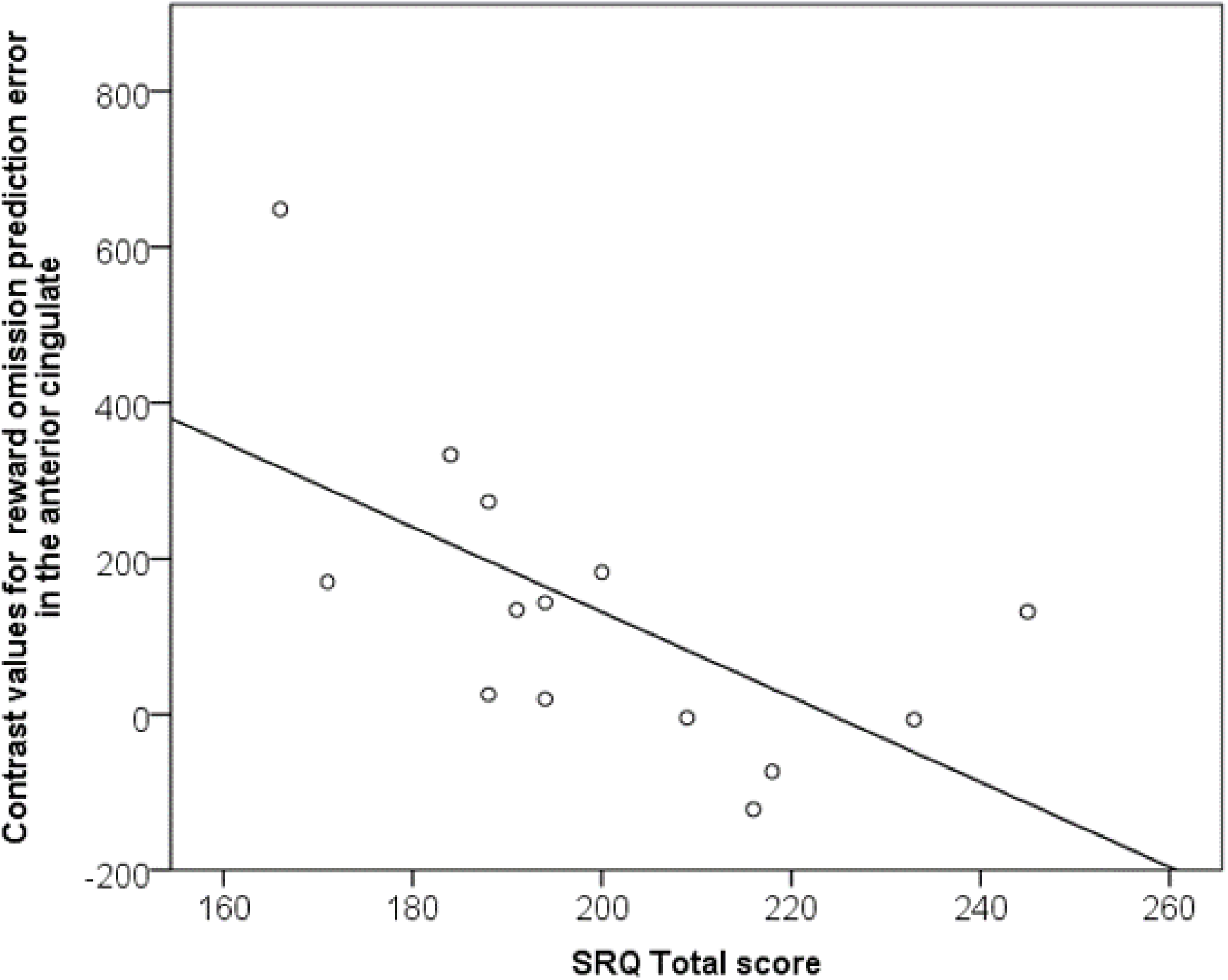
Significant correlation between self-regulation and negative prediction error activation in the anterior cingulate region of interest for OCD patients (rho=−0.7, p=0.002): stronger representation of prediction error during reward omission related to weaker self-regulation. SRQ – self-regulation questionnaire.

There were no correlations between placebo COPE values in the nucleus accumbens and rating scales (Y-BOCS, self-regulation including subscales, all p>0.5).

## Discussion

In this pharmacological fMRI study of reinforcement learning we found that OCD patients exhibited abnormally increased signaling of prediction errors to omitted rewards in the anterior cingulate cortex under placebo treatment, which was reduced by acute dopaminergic drug treatment using either pramipexole or amisulpride. This bidirectional reduction was unexpected and requires further analysis in future research. Furthermore, we found that OCD patients also showed abnormally increased activity associated with positive prediction errors in the nucleus accumbens in placebo, which was, however, unaffected by dopaminergic drug treatment.

In OCD, during placebo, the anterior cingulate representation of negative prediction error (prediction error during omitted rewards) was significantly greater than in controls. In controls, we did not detect a clear cingulate prediction error signal during omitted rewards. It is possible that our paradigm was not sufficiently sensitive to detect a negative prediction error signal in the control group, although it was clearly sufficiently sensitive to detect the pronounced signal in OCD. The lack of clear prediction errors in controls may have resulted from our procedure containing only 40 reward pair stimuli trials; for comparison, other prior prediction error fMRI studies have included larger numbers of potentially rewarding trials, such as 80 (Murray *et al*, 2008) or 90 (Pessiglione *et al*, 2006). Pronounced anterior cingulate prediction error signals are consistent with a recent study reporting stronger fronto-striatal reward prediction error signals in OCD, also using a fMRI reinforcement learning paradigm (Hauser *et al*, 2017). Furthermore, exaggerated responses to errors is a consistent finding in OCD, with a number of prior studies showing an overactivation of frontal responses to error processing in OCD in both fMRI research (Stern *et al*, 2011; Ursu *et al*, 2003) and EEG Error Related Negativity studies (Gillan *et al*, 2017; Mathews *et al*, 2012).

One previous study of reward anticipation (not learning) showed nucleus accumbens dysfunction in OCD (Figee *et al*, 2011). Here, we report an overactivation in the nucleus accumbens responses to positive prediction errors in OCD, consistent with prior data showing overactivation in OCD across error tasks and brain regions (Stern *et al*, 2011; Ursu *et al*, 2003) and in line with results from Hauser and colleagues (Hauser *et al*, 2017), who showed excessive striatal reward prediction errors in OCD during reversal learning.

We found no significant correlations between anterior cingulate prediction error signals and a general OCD symptom score; this lack of association is in accordance with the results from the only other fMRI study of reward prediction error in OCD (Hauser *et al*, 2017). However, we did demonstrate an association between anterior cingulate negative prediction error signal with self-perceived ‘self-regulation’: driven by associations with subcomponents of difficulty in triggering behavioral change, searching for options, and implementing a plan. This result may be relevant to much recent data suggesting an impairment of goal-directed behavior in OCD, and highlights the importance of cingulate dysfunction in this impairment (Giele *et al*, 2016; Gillan and Robbins, 2014). Our data suggest that the self-regulation questionnaire may reflect underlying anterior cingulate function in OCD. We are not aware of any clinical trials for OCD that have used this scale as an outcome measure; however, our results suggest that including a questionnaire measure of goal directed behavioral control, such as the SRQ, might be useful in future clinical trials of interventions targeting cingulate mediated deficits in OCD.

Although the literature provides consistent evidence linking error processing to dopaminergic function (Schultz *et al*, 1997) – a system apparently impaired in OCD (Denys *et al*, 2004; Hesse *et al*, 2005; Moresco *et al*, 2007; Olver *et al*, 2009, 2010; Perani *et al*, 2008; Sesia *et al*, 2013; Van Der Wee *et al*, 2004) – our study extends prior work by directly showing the effect of dopaminergic modulation on brain prediction error responses in OCD. Over-activation of the anterior cingulate cortex in response to negative reward prediction errors in OCD was remediated by amisulpride, the selective D_2/3_ receptor antagonist, as hypothesized. However, the D2/3 agonist pramipexole unexpectedly had similar, rather than opposite, effects.

Nevertheless, previous studies have shown that both drugs can exert bidirectional effects on reward prediction error associated responses. Thus, amisulpride or similar D_2/3_ receptor antagonists can either enhance (Jocham *et al*, 2011) or depress (Abler *et al*, 2007) reward related brain activity. Furthermore, dopamine agonist-like medications have also been shown to be capable of enhancing (Pessiglione *et al*, 2006) or attenuating (Bernacer *et al*, 2013) reward prediction error associated responses, with the directionality of the effect being dependent on the particular task, agonist and dose employed. Two obvious accounts of this pattern of effects are that either (I) both agents in our study reduce prediction errors by operating at opposite sides of a hypothetical inverted U-shaped function that determines the net effect of dopaminergic modulation on the computation of prediction error; or (II) that actions at inhibitory presynaptic receptors, at the level of the ventral tegmental area (Grace and Bunney, 1983) and/or via terminal autoreceptors (e.g. (Horst *et al*, 2019)), may be implicated for one or other drug that are opposite in sign to their post-synaptic effects. As the dosage of amisulpride in our study is between a high and a low dose, the latter account is further supported by the finding that low doses of amisulpride in animal studies are associated with a facilitation of behaviors depended on presynaptic dopamine receptor activation, whereas higher doses were found to decrease behaviors associated with postsynaptic receptor activation (Perrault *et al*, 1997). Such an action can only be assessed by determining an entire dose response curve, and it is an obvious limitation of the present data that this was not feasible in this study of human participants. In particular, it is possible that the low dose (0.5mg) of pramipexole employed in our study may have resulted primarily in presynaptic effects at presynaptic D2 autoreceptors which have higher affinity for dopamine than postsynaptic heteroreceptors (Ford, 2014). It is also possible that the degree to which our study drugs act pre-versus post-synaptically may depend on dopamine receptor availability, presynaptic action being favored when receptor availability is low. We note prior evidence of low dopamine receptor availability in OCD (Denys *et al*, 2004), which is likely to contribute to differential drug effects between OCD patients and controls and is consistent with the observed effects of these dopaminergic agents. We initially intended to use a higher dose – 1.5mg daily –the lower end of the target daily dose in Parkinson’s Disease (Constantinescu, 2008). However, in clinical practice it is advised to titrate up to this dose and indeed on initial testing we found that acute administration of 1.5mg without prior titration was poorly tolerated, necessitating our decision to use the lower dose of 0.5mg. There are two further limitations, the first is that 16 out of the 18 patients were taking medication for OCD at the time of the study, though the dopaminergic drug treatment effect was present in OCD irrespective of SSRI treatment (high vs low dose). A second possible limitation is the relatively modest sample sizes. The study may have been underpowered to determine clear differences for some group comparisons. A retrospective power analysis however provided reasonable confidence levels for correctly rejecting the null hypothesis (1-B>0.7).

In some circumstances pramipexole impairs behavioral learning, possibly by disrupting phasic dopaminergic signaling, in reinforcement learning tasks (Nagy *et al*, 2012; Pizzagalli *et al*, 2008), and other learning or decision making tasks (Gallant *et al*, 2016). In stimulant dependent individuals, pramipexole has also been shown to reduce behavioral perseveration during learning tasks (Ersche *et al*, 2011), a behavior frequently found in OCD (Giele *et al*, 2016; Hauser *et al*, 2017; Morris *et al*, 2017). However, Ersche et al (2011) found no evidence for normalization of behavioral perseveration in the same set of OCD patients as used in this study. We also observed no effects of dopaminergic drugs on our behavioral choice measures of learning in OCD patients or healthy controls although pramipexole did lengthen response times on reward trials, possibly consistent with previous findings reporting a slowing of reward learning after a single dose of pramipexole (Pizzagalli *et al*, 2008). In contrast, we did demonstrate that the fMRI reward prediction error signal in the anterior cingulate cortex was sensitive to dopaminergic manipulation in OCD, consistent with previous observations in a similar reinforcement learning task that fMRI measures of learning may be more sensitive than behavioral indices (Murray *et al*, 2010).

Although there have been no randomized controlled clinical effectiveness trials of amisulpride in OCD, meta-analysis confirms that dopamine receptor antagonists reduce obsessive-compulsive symptoms in SSRI-refractory OCD (Dold *et al*, 2015; Veale *et al*, 2014). Whilst the addition of an dopamine receptor antagonist medication is a commonly used augmentation strategy for patients who do not recover completely on first line treatments (Hirschtritt *et al*, 2017), the mechanisms through which this leads to clinical improvement are not fully understood. Our data suggest one possible mechanism – remediation of otherwise excessive anterior cingulate error signaling in OCD that are related to the self-regulatory impairments we found in these patients. In contrast, we found no effect of dopaminergic drug treatment on the enhanced positive reward prediction error signals expressed in the nucleus accumbens of these OCD patients, which was not shown however to be related to symptoms in this patient sample.

There have been no clinical trials of pramipexole for OCD, although three small clinical trials have shown evidence that the related drug, dexamphetamine, can be effective in OCD (Insel *et al*, 1983; Joffe *et al*, 1991; Koran *et al*, 2009). Our results may prompt consideration of further research into low-dose D_2/3_ agonist treatment for OCD, although here considerable caution, rigorous risk assessment and very careful experimental design is advised given prior reports that pramipexole may cause or worsen compulsive behaviors in some people (Kolla *et al*, 2010). Our data are consistent with the potential therapeutic actions of partial D_2/3_ agonists; one such drug, aripiprazole, has already been shown to be effective in augmentation in OCD, with some evidence of efficacy at very low doses (Ercan *et al*, 2015; Janardhan *et al*, 2017; Muscatello *et al*, 2011; Sayyah *et al*, 2012; Veale *et al*, 2014). Additional mechanistic and pragmatic research is warranted to determine which OCD patients are most likely to benefit from dopaminergic treatment.

In summary, these data extend previous evidence of abnormal processing of errors in the fronto-striatal networks in OCD, by showing that the heightened sensitivity of anterior cingulate cortex in OCD can be reduced by dopaminergic modulation. This study therefore is the first to provide direct evidence of dopaminergic dysfunction during reward prediction error signaling in OCD.

## Acknowledgments

Supported by an award from GSK to University of Cambridge (RG45422, Principal Investigator TWR), a Wellcome Trust Senior Investigator Award (104631/Z/14/Z) to TWR, by a MRC Clinician Scientist award (G0701911) to GKM, and by a Betty Behrens Research Fellowship of Clare Hall to KDE. The research was conducted in the University of Cambridge Behavioural and Clinical Neuroscience Institute which was supported by a joint award from the Medical Research Council (G1000183) and Wellcome Trust (093875/Z/10Z)

