## Supplementary materials for "Dopaminergic drug treatment remediates exaggerated cingulate prediction error responses in obsessive-compulsive disorder"

*fMRI task*

During the fMRI scan, subjects carried out an instrumental discrimination learning task that required making choices to maximize wins and minimize losses (Figure 1). In each trial, one of three possible pairs of abstract pictures was randomly presented: rewarding, punishing or neutral. There were 40 trials of each valence (120 trials per visit in total), randomized and interspersed by an inter-trial interval of variable duration (0.5-4.5 s). For each trial, the subject used a button push to indicate a choice of picture. Outcomes on trials where the potentially rewarding pair were shown could be a win (a picture of a 50 pence coin in rewarding trials with the text “you won 50p”) or no feedback. Outcomes on trials with the potentially punishing pair were a loss (a red cross over the same picture of the coin in punishing trials with the text “you lost 50p”) or no feedback. On neutral trials, there was neutral (“no change”) feedback or no feedback. Each pair of stimuli had one stimulus that, when chose, was associated with one of the two potential outcomes with 70% probability, and the other potential outcome with 30% probability. Subjects learned the task by trial and error. Optimal responding involved learning to choose the high probability and the low probability cues in rewarding and punishing trials, respectively. The relationship of a given abstract picture to the probability of obtaining the expected feedback was counterbalanced across subjects. The selection of the cue was by pressing one of two buttons (left or right), with the first two fingers of the right hand. The position of the high probability stimulus was also randomized and counterbalanced across trials within valence. Before starting the task, subjects were informed that any money they won during the task would be paid to them at the end of the experiment. We focus in this report on how participants learned about rewards; hence we report fMRI results from the reward trials only.

*MRI acquisition*

Whole-brain MRI data were acquired at the Wolfson Brain Imaging Centre, University of Cambridge, Cambridge, UK, using a Siemens Magnetom Tim Trio whole-body scanner operating at 3 T (Siemens Medical Solutions, Erlangen, Germany). During the performance of the learning task, 32 transaxial sections of gradient echo, echoplanar imaging (EPI) data depicting blood oxygen level–dependent contrast were acquired parallel to the intercommissural line with the following parameters: repetition time=2000 milliseconds, echo time=30 milliseconds, flip angle = 78°, slice thickness = 3 mm plus 0.75-mm interslice gap, image matrix size=64 X 64, and within-plane voxel dimensions = 3.0 mm X 3.0 mm. 404 volumes were collected per visit. Prior to data analysis, the first 5 echoplanar images were discarded to account for T1 equilibration effects. A 3D MPRAGE structural T1 scan was also acquired on each participant to facilitate standard space mapping.

*Computational model*

We fitted a standard reinforcement learning algorithm to each subject’s sequence of choices: a Q learning algorithm, which has been shown previously to offer a good account of instrumental choice in humans and other primates. For each pair of stimuli, A and B, the model estimates the expected values of choosing A(Qa) and choosing B(Qb), on the basis of individual sequences of choices and outcomes. This value, termed a Q value, is essentially the expected reward obtained by taking that particular action. These Q values were set at zero before learning, and after every trial t>0 the value of the chosen stimulus (say A) was updated according to the rule

Q_A_(t+1) = Q_A_(t) + α * δ(t)

The prediction error was

δ (t) = R(t) – Q(t)

where R(t) is defined as the reinforcement obtained as an outcome of choosing A at trial t. In other words, the prediction error d (t) is the difference between the expected outcome (that is, Q(t)) and the actual outcome (that is, R(t)). The reinforcement magnitude R was 1 for feedback and 0 for ‘nothing’ outcomes. Given the Q values, the associated probability of selecting each action was estimated by implementing the softmax rule, for example, for choosing A,

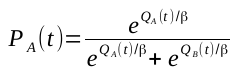

This is a standard stochastic decision rule that calculates the probability of taking one of a set of actions according to their associated values. The constants α (learning rate) and β (noise or exploration-exploitation parameter) were adjusted to maximize the probability (or likelihood) of the actual choices under the model. To compare the accuracy of fit between diagnoses and conditions, we used negative log likelihood. We estimated the model parameters α and β for each individual participant, and used these to calculate vectors of estimated prediction errors (one prediction error estimate per trial).

*SSRI medication*

16 of 18 patients were taking SSRI medication. We defined high dose as fluoxetine 60mg or above; citalopram 60 mg or above; sertraline 200mg or above; paroxetine 50mg or above; escitalopram 20mg or above; fluvoxamine 200mg daily or above. Doses below this were operationalized as low dose. This split revealed 9 patients on high dose SSRIs and 9 patients on low dose or no medication.

*Behavioral results – Punishment and neutral trials*

*Discrimination learning:* For punishment trials, we found a significant diagnosis effect (F=17.0, dfd=1, dfe= 34, p<0.001, partial ƞ^2^=0.33), where controls performed better than patients. There was no effect of drug (p=0.66) or drug X diagnosis interaction (p=0.31). For neutral trials, we did not find any significant effect (diagnosis: p=0.84; drug: p=0.47; drug X diagnosis: p=0.43). (See Supplementary Figure 2.)

*Reaction times:* Neither for punishment trials, nor for neutral trials, we found any significant differences (Punishment: diagnosis: p=0.75; drug: p=0.44; drug X diagnosis: p=0.36; Neutral: diagnosis: p=0.32; drug: p=0.38; drug X diagnosis: p=0.22).

*Win-stay/Lose-shift behavior – all trial types*

Furthermore, we analyzed the win-stay/lose-shift behavior across all trial types, to examine in more detail whether decision-making was influenced by drug treatment. We took into account all decisions made and analyzed how often participants chose the same stimulus that led to an optimal outcome on a subsequent trial of a particular condition (win-stay), and how often they altered their decision after a loss (lose-shift). When analyzing the win-stay/lose-shift behavior we did not find any significant differences with regard to diagnosis or drug treatment, or interaction. However, we found a significant effect for trial type (F=133.6, dfd=2, dfe= 34, p<0.001, partial ƞ^2^=0.80), indicating best performance in reward trials and worst performance in neutral trials. Furthermore we found significant effect for win-stay/lose-shift- behavior (F=296.6, dfd=1, dfe= 34, p<0.001, partial ƞ^2^=0.90), showing a high tendency to repeat a decision after a win, and a low tendency to shift after a loss. (See Supplementary Table 2).

*Voxelwise analysis, pooled data (“main effect of task”)*

We analyzed voxelwise fMRI data using permutation testing with the FSL tool randomize and threshold free cluster enhancement (Smith & Nichols, 2009), collapsing over drug conditions, and pooling patients and controls. The analysis was conducted in a ROI encompassing entire frontal cortex and striatum, family-wise error corrected for multiple comparisons (p<0.05). On voxelwise analysis, positive reward prediction error was associated with activation in the striatum (including nucleus accumbens), ventromedial prefrontal cortex and anterior cingulate cortex, and negative reward prediction error was associated with activation in the frontal pole, left middle frontal gyrus and left postcentral gyrus.

| **Supplementary Table 1.** Choice Performance and Reaction Times for reward trials across all treatments | | | | | | | |
| --- | --- | --- | --- | --- | --- | --- | --- |
|  |  |  |  | **N. of “correct” choices** | | **Reaction times (ms)** | |
| *Drug* | *Valence* | *Group* |  | *Mean (± SD)* | | *Mean (± SD)* | |
| *Placebo* | *Reward* | *Controls* |  | 34.83 | (9.82) | 766.29 | (161.7) |
|  |  | *OCD* |  | 28.89 | (14.56) | 775.57 | (118.9) |
| *Pramipexole* | *Reward* | *Controls* |  | 30.28 | (12.33) | 843.02 | (161.0) |
|  |  | *OCD* |  | 27.56 | (14.86) | 847.56 | (188.6) |
| *Amisulpride* | *Reward* | *Controls* |  | 35.17 | (9.46) | 733.82 | (140.0) |
|  |  | *OCD* |  | 30.50 | (12.35) | 781.90 | (175.0) |

| **Supplementary Table 2.** Win-stay/Lose-shift behavior for all trial types across all treatments. | | | | | | | |
| --- | --- | --- | --- | --- | --- | --- | --- |
|  |  | **Placebo** | | **Pramipexole** | | **Amisulpride** | |
| Reward |  | Mean | SD | Mean | SD | Mean | SD |
| Win-stay | Control | 97.5 | 5.2 | 91.0 | 16.9 | 97.0 | 5.0 |
|  | OCD | 93.6 | 10.7 | 91.0 | 13.6 | 93.6 | 7.8 |
| Lose-shift | Control | 11.3 | 13.3 | 19.6 | 18.8 | 12.1 | 16.3 |
|  | OCD | 12.9 | 12.6 | 15.4 | 16.3 | 19.0 | 18.1 |
| Punishment | | Mean | SD | Mean | SD | Mean | SD |
| Win-stay | Control | 88.2 | 9.5 | 81.5 | 14.1 | 92.2 | 8.9 |
|  | OCD | 84.3 | 10.8 | 80.2 | 15.3 | 81.7 | 16.3 |
| Lose-shift | Control | 55.6 | 17.3 | 58.9 | 15.4 | 59.6 | 18.8 |
|  | OCD | 63.1 | 17.4 | 59.0 | 12.3 | 58.8 | 14.6 |
| Neutral |  | Mean | SD | Mean | SD | Mean | SD |
| Win-stay | Control | 63.7 | 24.32 | 65.0 | 24.2 | 65.9 | 18.4 |
|  | OCD | 70.7 | 24.3 | 67.2 | 21.5 | 59.2 | 20.9 |
| Lose-shift | Control | 27.5 | 18.1 | 27.7 | 18.7 | 29.8 | 17.6 |
|  | OCD | 23.2 | 22.5 | 26.3 | 17.2 | 33.9 | 19.6 |

A Learning Rate on Reward Trials

Proportion of correct choices (%)

| Number of Trials | Controls Placebo | Controls Amisulpride | Controls Pramipexole | OCD Placebo | OCD Amisulpride | OCD Pramipexole |
| --- | --- | --- | --- | --- | --- | --- |
| 1 | 55.5556 | 55.5556 | 44.4444 | 55.5556 | 50 | 44.4444 |
|  | 77.7778 | 77.7778 | 55.5556 | 50 | 55.5556 | 50 |
|  | 83.3333 | 72.2222 | 72.2222 | 55.5556 | 66.6667 | 55.5556 |
|  | 83.3333 | 83.3333 | 72.2222 | 61.1111 | 50 | 55.5556 |
| 5 | 72.2222 | 77.7778 | 77.7778 | 61.1111 | 61.1111 | 61.1111 |
|  | 83.3333 | 88.8889 | 66.6667 | 61.1111 | 83.3333 | 61.1111 |
|  | 88.8889 | 88.8889 | 55.5556 | 66.6667 | 50 | 72.2222 |
|  | 94.4444 | 83.3333 | 66.6667 | 72.2222 | 66.6667 | 72.2222 |
|  | 94.4444 | 88.8889 | 72.2222 | 72.2222 | 72.2222 | 61.1111 |
| 10 | 94.4444 | 77.7778 | 66.6667 | 72.2222 | 72.2222 | 72.2222 |
|  | 94.4444 | 88.8889 | 72.2222 | 66.6667 | 66.6667 | 66.6667 |
|  | 88.8889 | 83.3333 | 66.6667 | 61.1111 | 72.2222 | 66.6667 |
|  | 94.4444 | 88.8889 | 66.6667 | 66.6667 | 72.2222 | 66.6667 |
|  | 88.8889 | 88.8889 | 66.6667 | 77.7778 | 72.2222 | 77.7778 |
| 15 | 88.8889 | 88.8889 | 72.2222 | 83.3333 | 77.7778 | 72.2222 |
|  | 88.8889 | 88.8889 | 77.7778 | 72.2222 | 77.7778 | 72.2222 |
|  | 88.8889 | 88.8889 | 77.7778 | 72.2222 | 77.7778 | 66.6667 |
|  | 88.8889 | 88.8889 | 77.7778 | 77.7778 | 77.7778 | 61.1111 |
|  | 88.8889 | 88.8889 | 72.2222 | 66.6667 | 88.8889 | 66.6667 |
| 20 | 83.3333 | 88.8889 | 77.7778 | 77.7778 | 72.2222 | 77.7778 |
|  | 77.7778 | 88.8889 | 77.7778 | 77.7778 | 77.7778 | 66.6667 |
|  | 83.3333 | 94.4444 | 83.3333 | 77.7778 | 83.3333 | 66.6667 |
|  | 83.3333 | 94.4444 | 83.3333 | 72.2222 | 72.2222 | 66.6667 |
|  | 83.3333 | 83.3333 | 83.3333 | 72.2222 | 77.7778 | 72.2222 |
| 25 | 77.7778 | 94.4444 | 83.3333 | 72.2222 | 83.3333 | 66.6667 |
|  | 88.8889 | 88.8889 | 77.7778 | 77.7778 | 77.7778 | 72.2222 |
|  | 88.8889 | 88.8889 | 83.3333 | 77.7778 | 77.7778 | 72.2222 |
|  | 83.3333 | 94.4444 | 83.3333 | 77.7778 | 77.7778 | 77.7778 |
|  | 88.8889 | 94.4444 | 88.8889 | 72.2222 | 83.3333 | 72.2222 |
| 30 | 88.8889 | 88.8889 | 83.3333 | 77.7778 | 88.8889 | 77.7778 |
|  | 88.8889 | 94.4444 | 83.3333 | 77.7778 | 88.8889 | 83.3333 |
|  | 88.8889 | 83.3333 | 83.3333 | 77.7778 | 88.8889 | 83.3333 |
|  | 94.4444 | 94.4444 | 83.3333 | 77.7778 | 88.8889 | 72.2222 |
|  | 94.4444 | 94.4444 | 83.3333 | 72.2222 | 83.3333 | 72.2222 |
| 35 | 88.8889 | 88.8889 | 72.2222 | 77.7778 | 83.3333 | 83.3333 |
|  | 88.8889 | 94.4444 | 88.8889 | 72.2222 | 88.8889 | 66.6667 |
|  | 94.4444 | 94.4444 | 83.3333 | 83.3333 | 83.3333 | 77.7778 |
|  | 94.4444 | 94.4444 | 88.8889 | 77.7778 | 88.8889 | 66.6667 |
|  | 94.4444 | 94.4444 | 77.7778 | 83.3333 | 83.3333 | 72.2222 |
| 40 | 88.8889 | 94.4444 | 77.7778 | 83.3333 | 88.8889 | 66.6667 |

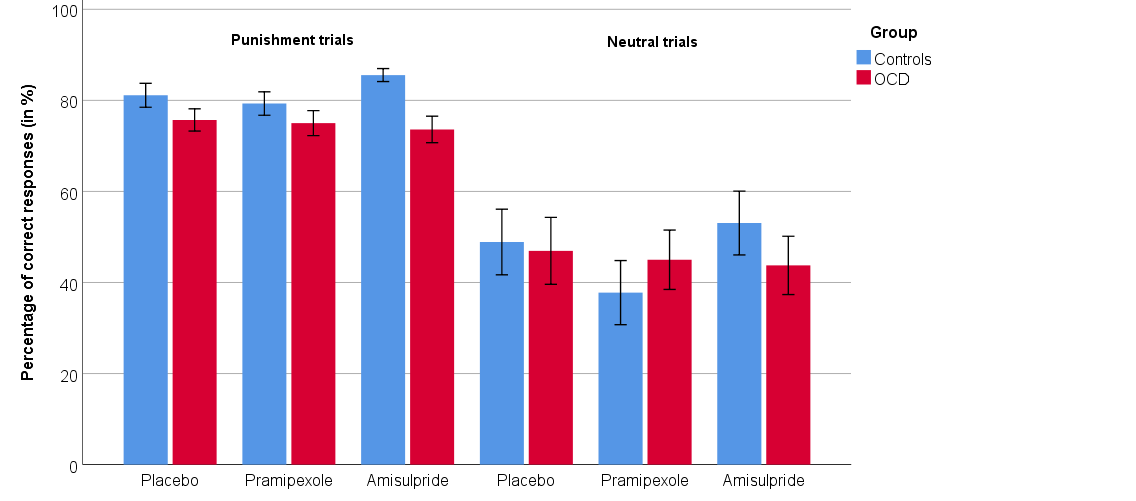
**Supplementary Figure 1.** The learning curves show a trial-by-trial depiction of observed correct “choices” per group (controls, OCD) and treatment (placebo, amisulpride, pramipexole). The ‘correct’ stimulus is associated with a probability of 70% of winning 50 pence.

**Supplementary Figure 2.** Choice performance (mean ±1 SEM) during punishment and neutral trials under different drugs, separated by group.

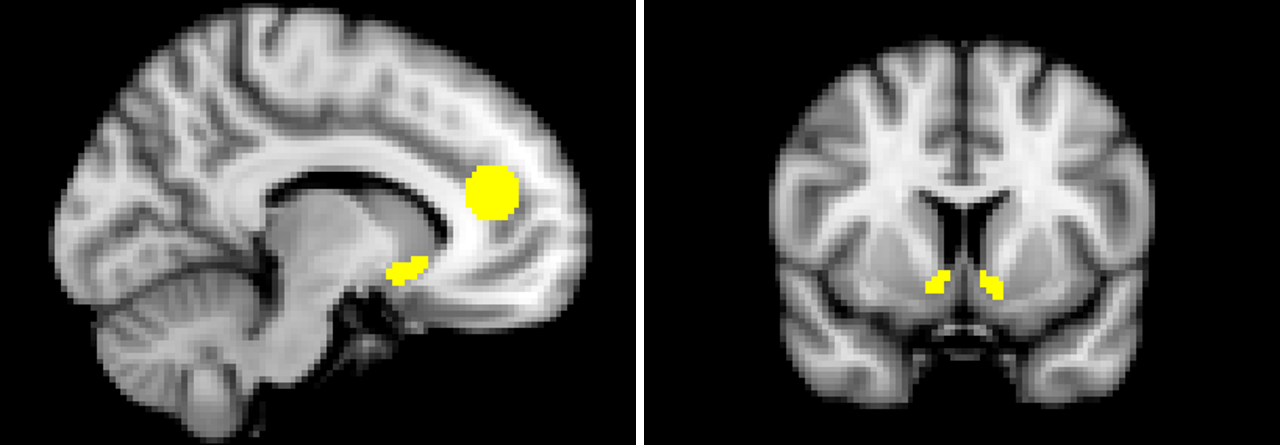

**Supplementary Figure 3.** Anatomical masks of regions of interest used for group by drug analysis. Areas marked in yellow show anatomical masks. (ACC: 1419 voxels, extracted utilizing a sphere, 14 mm, centered at x=0, y=42, z=18, size: 11.35 cm^3^, based on previous work (Hauser et al., 2017); full nucleus accumbens: 70 (left) and 56 (right) voxels, size: 0.56 (left) and 0.45 (right) cm^3^, extracted using Harvard-Oxford-subcortical atlas supplied with FSL.

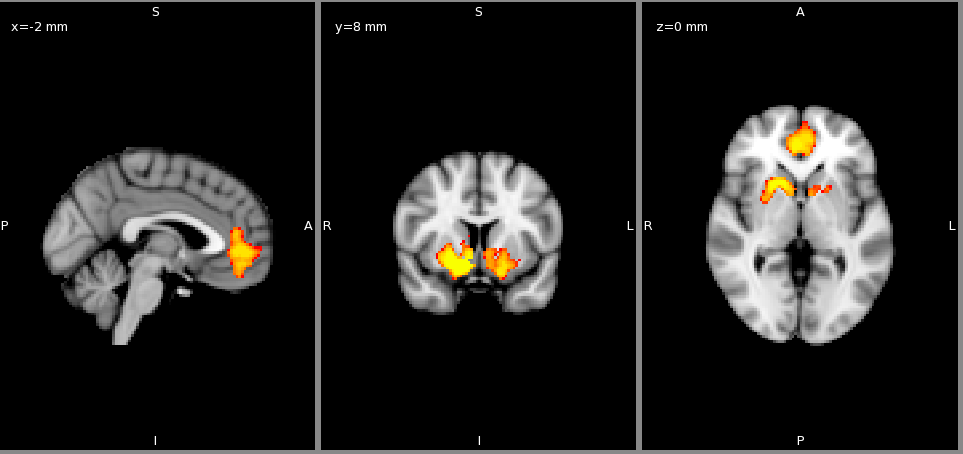

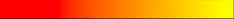

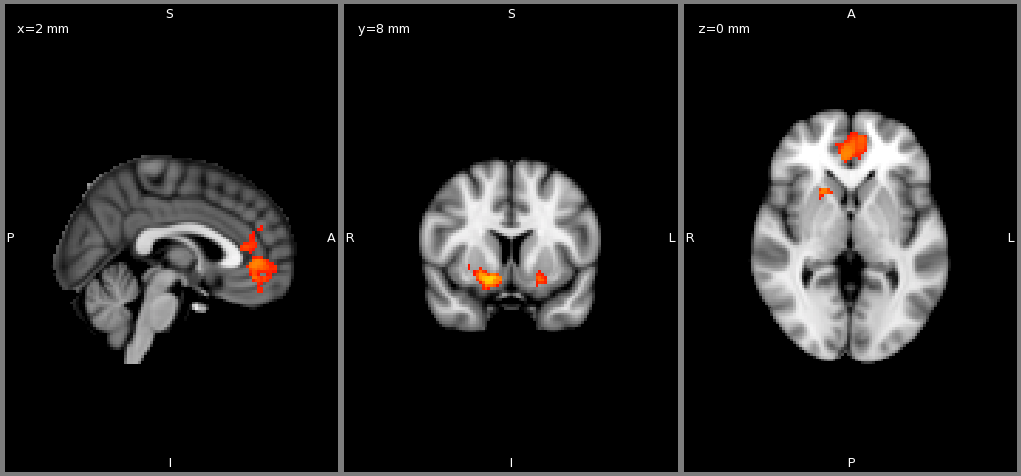
**Supplementary Figure 4.** Colored areas show voxels in entire sample (upper panel) and controls only (lower panel) whose activation correlates with positive reward prediction error (p<0.05 family wise error corrected for multiple comparisons across frontal cortex and striatum using randomize and threshold free cluster enhancement, 1000 permutations). Color coding ranges from yellow p=0.001 corrected to red p<0.05 corrected.

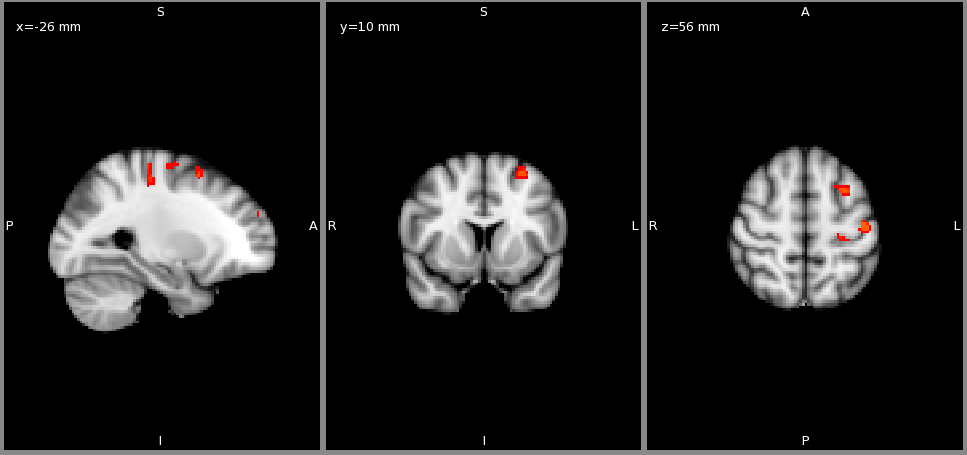

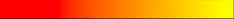
**Supplementary Figure 5.** Areas marked in yellow/red show voxels in entire sample whose activation correlates with negative reward prediction error (p<0.05 family wise error corrected for multiple comparisons across frontal cortex and striatum using randomize and threshold free cluster enhancement, 1000 permutations). Color coding ranges from yellow p=0.001 corrected to red p<0.05 corrected.
